# Reproduction and preference to macronutrients have different relations to biological or chronological age in *Drosophila*

**DOI:** 10.1101/2024.12.15.628509

**Authors:** Oleh Lushchak, Olha Strilbytska, Pavlo Petakh, Oleksandr Kamyshnyi, Oleksandr Koliada, Uliana Semaniuk

## Abstract

Genetic manipulations, dietary composition and supplementations with varied drugs and natural compounds were shown to extend both the life- and health-span in different model organisms. An understanding of the mechanisms behind the beneficial properties of intervention includes the evaluation of physiological and molecular traits at certain time points that reflect the values at distinguished chronological age. Thus, if one cohort has a long-lived phenotype than measurements at certain time points represent the difference between organisms of different biological ages. We have compared reproduction and consumption of specific macronutrients for flies of the same cohort but related to different quartiles of chronological and biological age. We found that the decline in carbohydrate or protein consumption was stronger in the case of chronological rather than biological age. However, flies of biological or chronological quartile 4 consumed virtually the same amounts of macronutrients. The decline in reproduction was significantly reduced in relation to biological age. Thus, the decline was about 38-68% when within chronological quartiles 2 and 1 but only 4-31% for biological ones. The reproductive capacity was reduced by 86-93% in flies of chronological Q4 as compared to a 60-77% decrease for those of biological. Starting from quartile 2 biologically aged flies laid significantly higher number of eggs than flies of the same chronological quartile. Our results point out the significant difference in flies of the same biological and chronological quartile and raise the question about the suitability of comparison traits of organisms with different lifespans same chronological age.

## 1 Introduction

In gerontological research distinguish between biological and chronological age of the studied objects. The chronological age is considered to be the elapsed time since birth. It is difficult to unify the term biological age. Biological age is an expression of typical phenomena of the normal course of life, id est age as such or age, which expresses the wear and tear of the organism and the actual physical condition of the assessed object. Biological age can be determined by biomarkers of aging, as well as by assessing the functional and structural changes in the body of the model object at a certain time of his chronological age. Biomarkers of aging are indicators that change with age according to certain patterns. Biological age can be faster or slower than chronological because everyone ages differently.

Biological age is often called functional or physiological (Mitnitski et al., 2002) and is an indicator of the general health status of an organism (Borkan and Norris, 1980). Moreover, biological age may help to predict the risk of age-related diseases or even mortality (Mitnitski et al., 2002). There are a set of aging biomarkers in humans including telomere length (Benetos et al., 2001), clotting factors and immune function tests (Seeman et al., 2001), glycosylated hemoglobin (Seeman et al., 2001), sarcopenia (Fisher, 2004) etc. The fruit fly *Drosophila melanogaster* has been used as a powerful animal model to study the physiology and genetics of aging (Heier et al., 2021). It was recently investigated parameters correlated with age and mortality in *Drosophila* that included negative geotaxis and walking speed, centrophobism behaviors, and pigments in oenocytes and eye (Tower, 2023). Indeed, walking speed and negative geotaxis decrease with age in flies of both sexes (Tower, 2023). Moreover, age-dependent functional deterioration and increased arrhythmia in *Drosophila* were shown (Blice-Baum et al., 2019). Intestinal barrier integrity might be used to assess biological age in *Drosophila* (Rera et al., 2012) and is associated with an increased risk of death. The study of Bushey and colleagues (2010) studied sleep characteristics as predictors of remaining lifespan in male *Drosophila*. Total body protein concentration was considered to be an important molecular marker of aging in *Drosophila* (Jacobson et al., 2010).

Experimental data with *Drosophila* showed that lifespan and aging are influenced by nutrition (Tatar et al., 2014; Strilbytska et al., 2024). The ratio of consumed protein relative to carbohydrates is among the certain determining factors related to longevity and metabolic traits (Strilbytska et al., 2020a; Strilbytska et al., 2022a). Dietary component concentration has a bell-shaped relationship with lifespan: poor dietary conditions lead to starvation or malnutrition, as well as enriched diets cause overfeeding and will induce a shortened lifespan (McCracken et al., 2020). Dietary restriction (DR) induces longevity in many species and maximizes lifespan (Tatar et al., 2014).

We aimed to test cohort survival, fecundity and feeding rate as reliable methods of biological age assessment, and to demonstrate the validity of these phenotypes for aging research. Studies of phenotypes of biological aging may provide novel strategies for beneficial lifestyle interventions for the alleviation of age-related functional decline.

## 2 Material and Methods

### 2.1 Fly husbandry

*Drosophila melanogaster* fruit flies of the *Canton-S* strain were used for this study. The flies were received from the Bloomington Stock Center (Indiana University, USA). Flies were kept at 25°C and 60-65% relative humidity, with overlapping generations and controlled density (Lozinsky et al., 2012), and were given standard food: 5% sucrose, 5% yeast, 6% corn, 1% agar, 0.18% methyl 4-hydroxyparabenzoic acid (methylparaben) and 0.6% propionic acid.

### 2.2 Experimental conditions

About 100 eggs were placed in 150-ml vials with 25 ml of food and allowed to develop. One-day-old flies were transferred into vials with fresh food without anesthesia and kept for an additional 4 days for mating. Before experiments, flies from multiple bottles were pooled. Female flies were placed individually into 7-ml vials with 1 ml of solidified water (1% agar-agar) and supplemented with two 5-μl microcapillary tubes (Drummond Scientific, Broomall, PA, USA) filled with liquid food. Four dietary regimens were prepared by paring 15% sucrose with 1.5 (1.5Y), 2.5 (2.5Y), 5 (5Y), or 15% (15Y) of autolyzed yeast. Foods were supplemented with 0.01% phosphoric and 0.1% propionic acid as antimould agents (Lee et al., 2008). Macronutrient consumption was calculated based on autolyzed yeast containing 45% protein, 24% carbohydrate (as glucose equivalents), 21% indigestible fibers, 8% water, and the remaining 2% fatty acids, minerals, and vitamins provided by the manufacturer (cat no. 103304; MP Biomedicals, Santa Ana, CA, USA).

### 2.3 Lifespan, Feeding and egg production

The mortality of flies was monitored daily. Capillaries were changed every other day and the volumes eaten were fixed. Three vials with capillary per condition were used as evaporation controls and mean values were subtracted from those eaten by fly (Koliada et al., 2020). Every other day flies were transferred into new vials and eggs were counted.

### 2.4 Data analysis

Feeding and fecundity were analyzed by 2-way ANOVA followed by Sidak multiple comparisons test to compare the effects related to different quarters and groups (Prism8; GraphPad Software, San Diego, CA, USA). All graphs were generated in «GraphpadPrism8».

## 3 Results

We have observed a lifespan of 52 days for most long-living flies in cohorts fed the diet with 1.5, 5, or 15% of yeast (Fig. 1). The lifespan of most long-living flies in a cohort fed by diet with 2.5% yeast concentration was 54 days. Thus, the chronological quartiles for flies of these groups were virtually the same. Shortage of biological quartile 1 (Q1) was observed when the concentration of yeasts was gradually increased from 1.5 to 15% (Fig. 1D). The duration of biological Q1 has shortened from 20 days at yeast concentration of 1.5% to 13 days at 15% yeast. Moreover, the shortage of Q1 was accompanied by an increased duration of Q4 that was almost twice longer for flies fed high-yeast diets (5 and 15% yeast) as compared to those restricted with yeast (1.5 and 2.5% yeast). These changes were observed because of different shapes of survival patterns with more flies dying at Q1 at higher concentrations of yeast in the diet.

**FIGURE 1.**
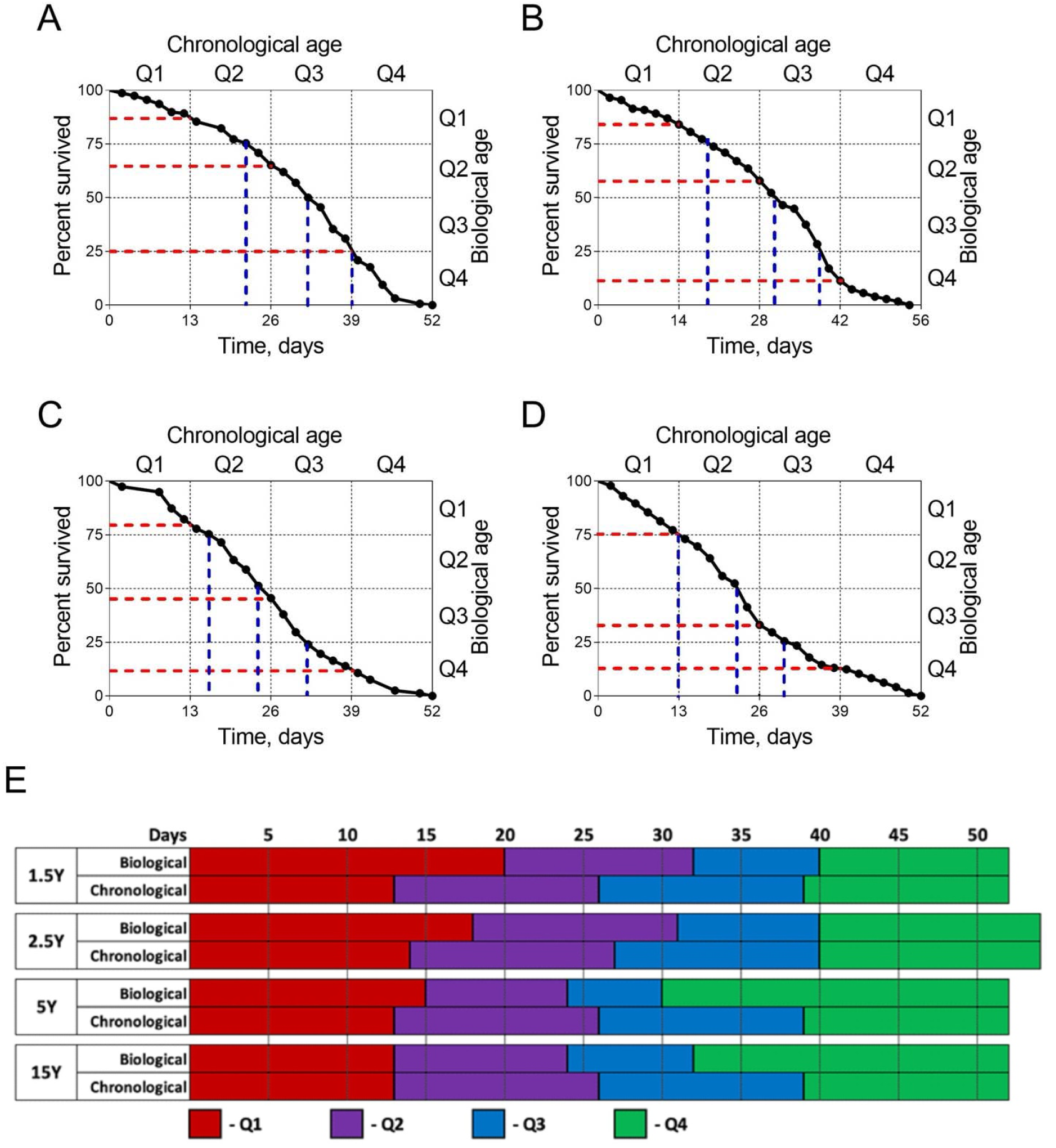
Lifespan and representation of quartiles for chronological and biological ages. Panels (**A**), (**B**), (**C**), and (**D**) represent the lifespan of flies at dietary regimes 1.5Y, 2.5Y, 5.0Y, and 15Y respectively. The red line demonstrates survival percent during the appropriate quartile of chronological age. The blue line demonstrates the relation of biological age to chronological age. (**E**) Representation of quartiles for chronological and biological ages.

Approximation of the curve crossing with every quartile of biological age on the X axis represents *Drosophila* biological age response landscapes by days and the relation to chronological age (*blue line*) (Fig. 1A-D). The Q1 of biological age refers to 75% of cohort survival and corresponds to 22 days of Q2 chronological age when flies consumed 1.5%Y food, 18 days of Q2 under 2.5Y food, 15 days under 5Y food, and 13 days of Q1 under 15Y diet. We defined the 50% survival cohort as Q2 of biological age. Female flies that consumed 1.5Y medium the Q2 of biological age matched up with 32 days of Q3 chronological age (Fig. 1A). Biological age of Q2 matched up with 32 days of Q3 chronological age when flies ate 2.5Y medium, 24 days of Q2 chronological age when flies were fed by 5Y and 15Y media (Fig. 1B-D). We also found the consistency of Q3 of the biological age which was defined as 25% survival with Q3 chronological age regarding the dietary protein (Fig. 1A). The Q3 of biological age corresponds to 39, 40, 26, and 23 days when females were reared on the medium 1.5Y, 2.5Y, 5Y, and 15Y respectively (Fig. 1B-D).

To compare dietary percent survival patterns, we approximate the curve crossing with every quartile of chronological age on the Y axis (*red line*) (Fig. 1A-D). At the end of Q1 of chronological age, the survival was 87, 85, 80, and 75% when the yeast concentration in the diet was 1.5, 2.5, 5, and 15% respectively. Similarly, the percent survival during Q2 of chronological age at these four diets was 65%, 60%, 45%, and ≈35%, respectively. Following an increase of yeast concentration in the diet from 1.5 to 15% the percent survival during Q3 decreased from 25% to 12% (Fig. 1A-D).

We next investigated how behavioral and physiological responses to yeast concentration in the diet differed between quartiles of chronological and biological age. The amount of carbohydrates consumed by female flies depends on the quartile and biological-chronological (BC-age) factor (Table 1). Females who consumed diets with different yeast concentrations ranging from 1.5% to 15% showed a decrease in carbohydrate intake from Q1 to Q4 within both biological and chronological ages (Fig. 2A-D). Moreover, we observed a 12% and 15% lower level of carbohydrate consumption in females of Q1 and Q2 chronological versus Q1 and Q2 biological age, respectively, under a 1.5Y diet (Fig. 2A; Sidak test: *p* < 0.04). Females fed by 2.5Y diet consumed the same amount of carbohydrates within Q1, Q2, and Q4 of biological and chronological age (Fig. 2B). However, during Q3 of chronological age females fed by 2.5Y diet consumed 21% less carbohydrates as compared to Q3 of biological age (Fig. 2B; Sidak test: *p* = 0.003). Lower level of carbohydrate consumption was observed in females fed by 5Y and 15Y diets during Q2 and Q3 of chronological age as compared to Q2 and Q3 of biological age by 30 and 23% respectively (Fig. 2C, D; Sidak test: *p* < 0.05).

**Table 3.**
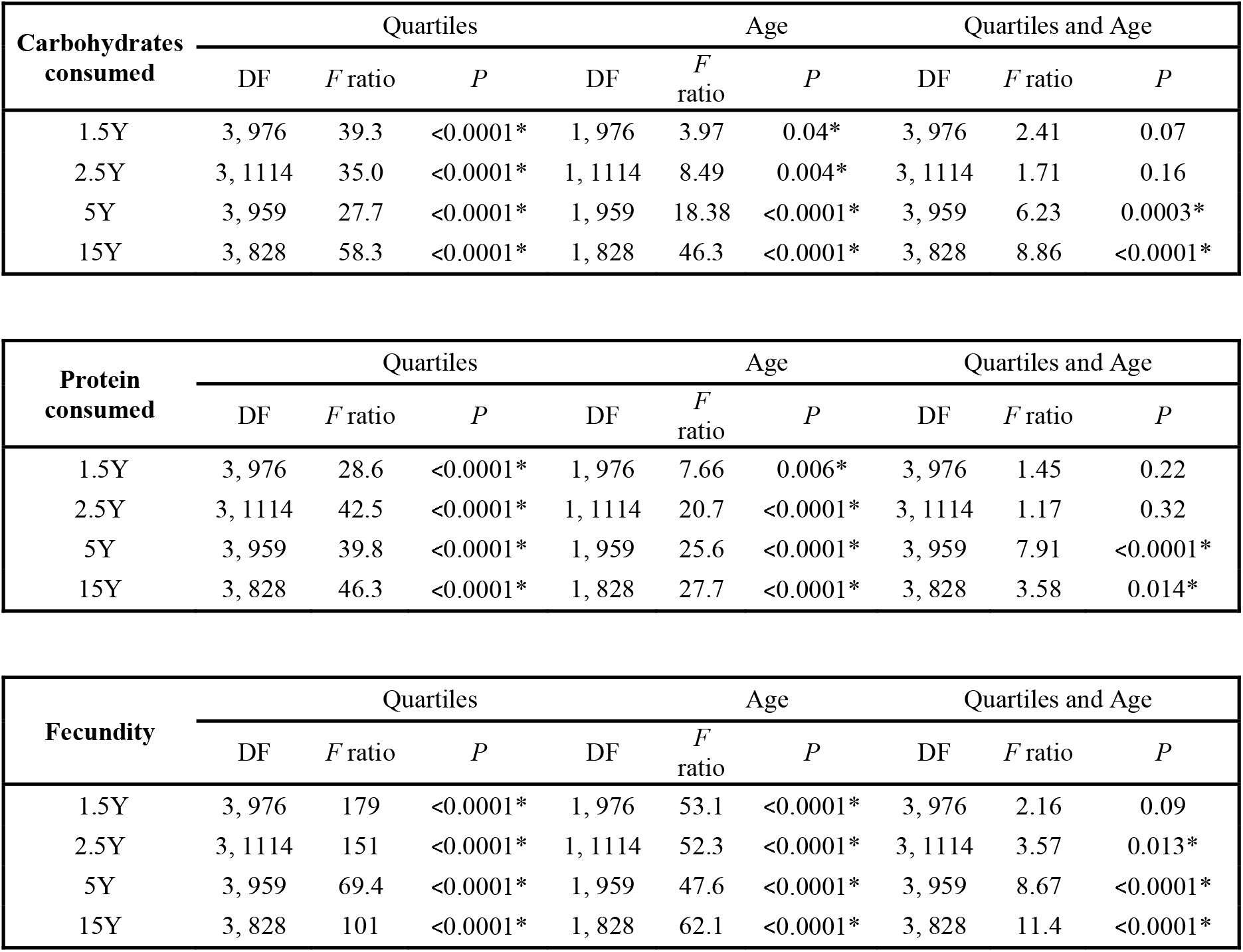
Two-way ANOVA to determine the impact of quartiles and BC age on parameters in parents that consumed diets with different autholyzed yeast concentration.

**FIGURE 2.**
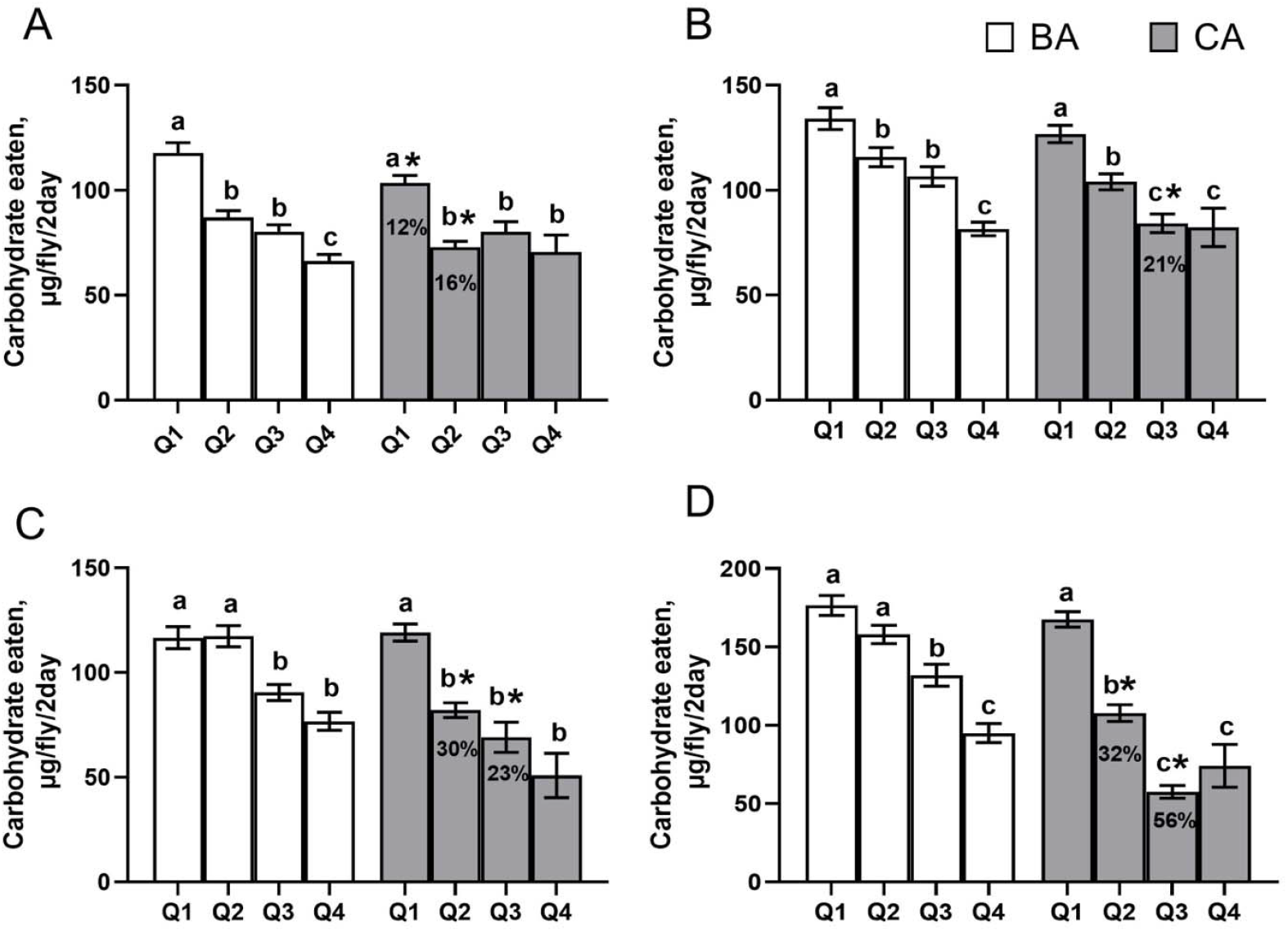
Amount of carbohydrate consumed during different quartiles of age. Flies were fed with food composed of 15% sucrose and 1.5% (**A**), 2.5% (**B**), 5.0% (**C**), and 15% (**D**) autolyzed yeast. Mean ± SEM; n = 142. *Significantly different from the values of the same quartile of biological age (Sidak’s multiple comparisons test, *p* < 0.05). The values were compared according to the Tukey test: a - indicates the highest average among all tested groups; b - indicates a significant difference from a with *p* <0.05; c is a significant difference from a and b with *p* < 0.05; d is a significant difference from a, b, and c with *p* < 0.05.

Both quartiles and BC-age factors had an impact on protein intake in females (Table 1). Protein consumption was the same during Q1 and Q4 of biological and chronological age (Fig. 3A-D). Lower protein intake was observed in female flies during Q2 of chronological versus biological age by 18-47% at all experimental dietary conditions (Fig. 3A-D; Sidak test: *p* < 0.05). Similarly, during Q3 of chronological age females consumed less protein on diets 2.5Y, 5Y, and 15Y as compared to Q3 of biological age (Fig. 3A-D; Sidak test: *p* < 0.05).

**FIGURE 3.**
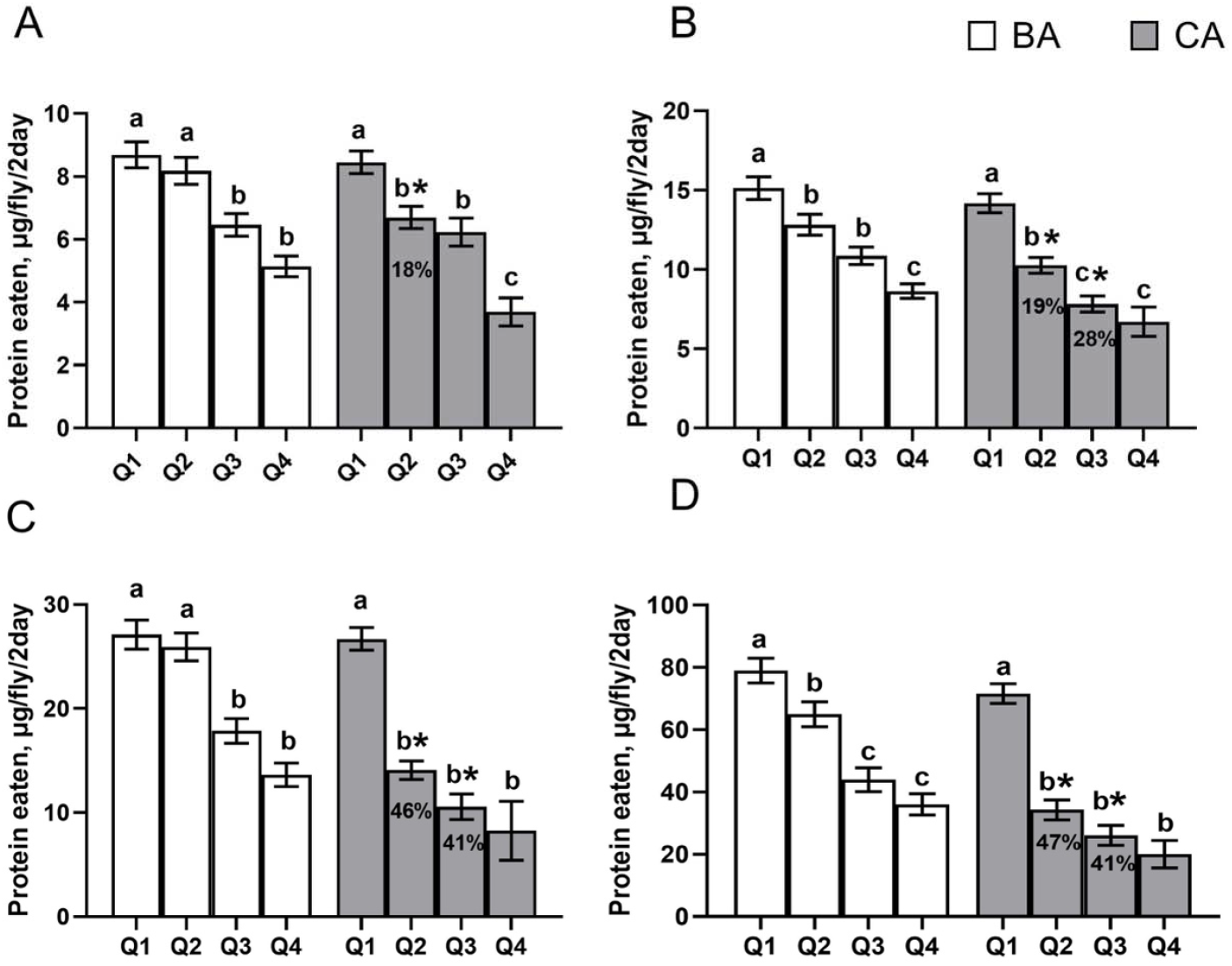
Amount of protein consumed during different quartiles of age. Flies were fed with food composed of 15% sucrose and 1.5% (**A**), 2.5% (**B**), 5.0% (**C**), and 15% (**D**) autolyzed yeast. Mean ± SEM; n = 142. *Significantly different from the values of the same quartile of biological age (Sidak’s multiple comparisons test, *p* < 0.05). The values were compared according to the Tukey test: a - indicates the highest average among all tested groups; b - indicates a significant difference from a with p <0.05; c is a significant difference from a and b with *p* < 0.05; d is a significant difference from a, b, and c with *p* < 0.05.

As expected, the highest fecundity rate was observed for females within Q1 and decreased significantly in Q4 regardless of the dietary regime (Fig. 4A-D). There was a difference in the egg-laying activity of females at the same quartile of biological versus chronological age. Lower levels of egg-laying activity were observed in females of Q1, Q2, Q3, and Q4 biological age as compared to appropriate quartiles of chronological age when flies were fed by 1.5Y medium by 13%, 42%, 50%, and 66%, respectively (Fig. 4A; Sidak test: *p* < 0.02). We did not find a significant difference in females’ fecundity rate during Q1 of biological versus chronological age under 2.5Y, 5Y, and 15Y dietary regimes (Fig. 4B-D). However, females of Q2, Q3, and Q4 biological age laid less eggs as compared to the same female cohorts within chronological age (Fig. 4B-D; Sidak test: *p* < 0.05).

**FIGURE 4.**
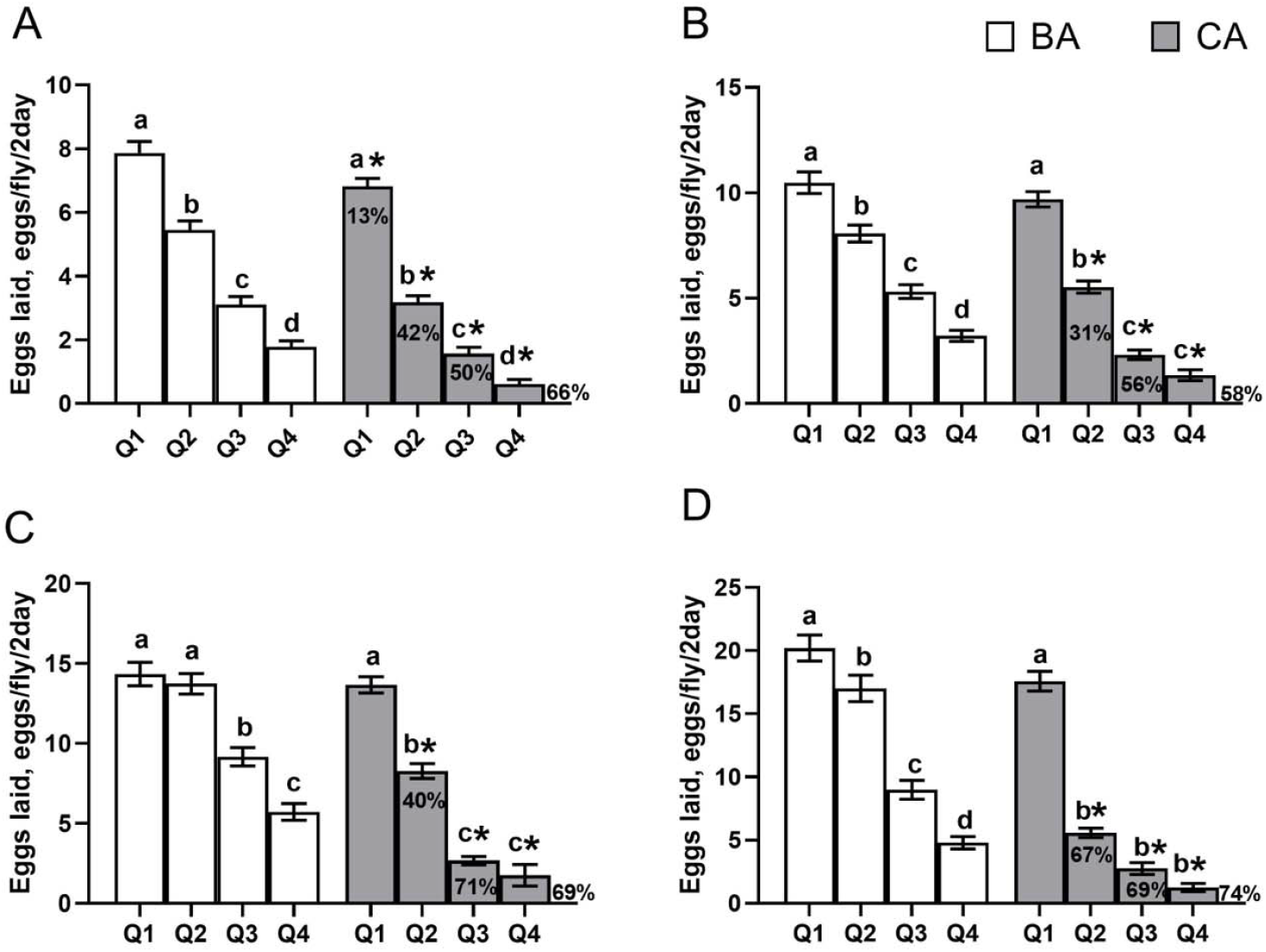
Amounts of eggs laid by flies as a function of age. Values are represented as mean ± SEM for 10-130 flies fed 15% sucrose and 1.5% (**A**), 2.5% (**B**), 5.0% (**C**), and 15% (**D**) autolyzed yeast. *Significantly different from the values of the same quartile of biological age (Sidak’s multiple comparisons test, *p* < 0.05). Bars sharing the same letter are not significantly different according to Tukey’s test.

## 4 Discussion

Validation of biomarkers of biological aging has been the focus of multiply gerontology research. Lifespan is the most robust measure of aging rate (Landis et al., 2020). Feeding and fecundity characteristics are related to age and are considered functional scoring systems in *Drosophila* for aging. Comprehensive functional assessment of feeding and fecundity improves our ability to understand whether dietary protein modulates fly longevity by acting on systemic aging or restricting the survival of the fly population.

We have previously shown that yeast concentration in the diet defines metabolism (Strilbytska et al., 2021), and antioxidant capacity and these factors are associated with lifespan (Strilbytska et al., 2021b; Strilbytska et al., 2022b). Hence, we try to explore the impact of dietary protein on the conformity between chronological and biological age. Differences in the rates of individual aging lead to differences between chronological and biological age observed in our studies. Flies within experimental cohorts aged at different rates. Our current study showed that yeast concentration in the diet significantly affects the conformity between chronological and biological age. We observed extended Q1 of biological age under consumption diets 1.5Y, 2.5Y, and 5Y. Females that were exposed to a 2.5Y diet showed a slightly prolonged lifespan as compared to the other experimental diets and the alignment of Q4 of biological and chronological aging. Prolonged Q4 within biological age in females exposed to 5Y and 15Y diets indicated the harmful effect of protein overconsumption (Lushchak et al., 2012; Strilbytska et al., 2021b). Previous studies showed that consumption of a diet with a high protein-to-carbohydrate ratio leads to lifespan shortening in *Drosophila* (Lushchak et al., 2012; Strilbytska et al., 2021b). Consumption of a high-protein diet could induce oxidative stress in fruit flies (Strilbytska et al., 2022b). Age-related changes in nutrient utilization in response to nutritional demands display *Drosophila* adaptation to systemic aging (Li et al., 2023). Dietary components determine the metabolism and feeding behavior of flies (Strilbytska et al., 2024). The metabolomic profiles of *Drosophila* were suggested to be a reliable biological clock and biomarker of aging (Zhao et al., 2020).

Determining age at death in a population does not provide comprehensive information about differences between young and old individuals and populations. Measuring some physiological elements of the aging process via analyzing any age-dependent physiological parameters might indicate the physiological age of the individual (Helfand and Rogina, 2003). Food intake is crucial to maintain energy homeostasis and link physiological needs to adaptive behaviors (Barnes et al., 2008). Nutrient consumption as well as the ratio of protein and carbohydrates consumed contribute significantly to longevity (Strilbytska et al., 2020a; Strilbytska et al., 2024). Within the current study, we found that both carbohydrate and protein intake are reduced in every next quartile of fly age. *Drosophila* insulin-like peptides (DILPs) are involved in the regulation of fly appetite (Semaniuk et al., 2021). The expression of *dilp3* in insulin-producing cells (IPCs) decreases with aging (Tanabe et al., 2017). Mutations of *dilp3* result in a greater appetite for specific dietary components (Semaniuk et al., 2021). Reduction in Akt activation and up-regulation of dFOXO target genes during aging confirm the involvement of insulin/insulin-like growth factor signaling (IIS) in the pathophysiology of aging (Rera et al., 2012; Strilbytska et al., 2020b). We also suggest that a reduction in the overall feeding rate might be associated with a decrease in gut functioning in aged flies. Indeed, it was previously shown progressive loss of barrier function in the aging *Drosophila* gut (Rera et al., 2012). Aged gut is characterized by significant changes in the composition, abundance, and function of the gut microbiota in fruit flies (Arias-Rojas and Iatsenko, 2022). Disruption of gut integrity predicts the death of individual flies regardless of chronological age (Rera et al., 2012). Loss of tissue integrity in aged flies is associated with increased antimicrobial peptide expression during aging (Rera et al., 2012).

Diet and macronutrient consumption are critical in shaping lifespan and healthspan (Strilbytska et al., 2024). Carbohydrate and protein consumption differed significantly during Q3 and Q4 of biological versus chronological age (Fig. 5). Protein restriction causes a significant difference in carbohydrate intake during Q1 of biological versus chronological age (Fig. 5A). We also found higher macronutrient consumption within quartiles of biological as compared to the same quartiles of chronological age (Fig. 5).

**FIGURE 5.**
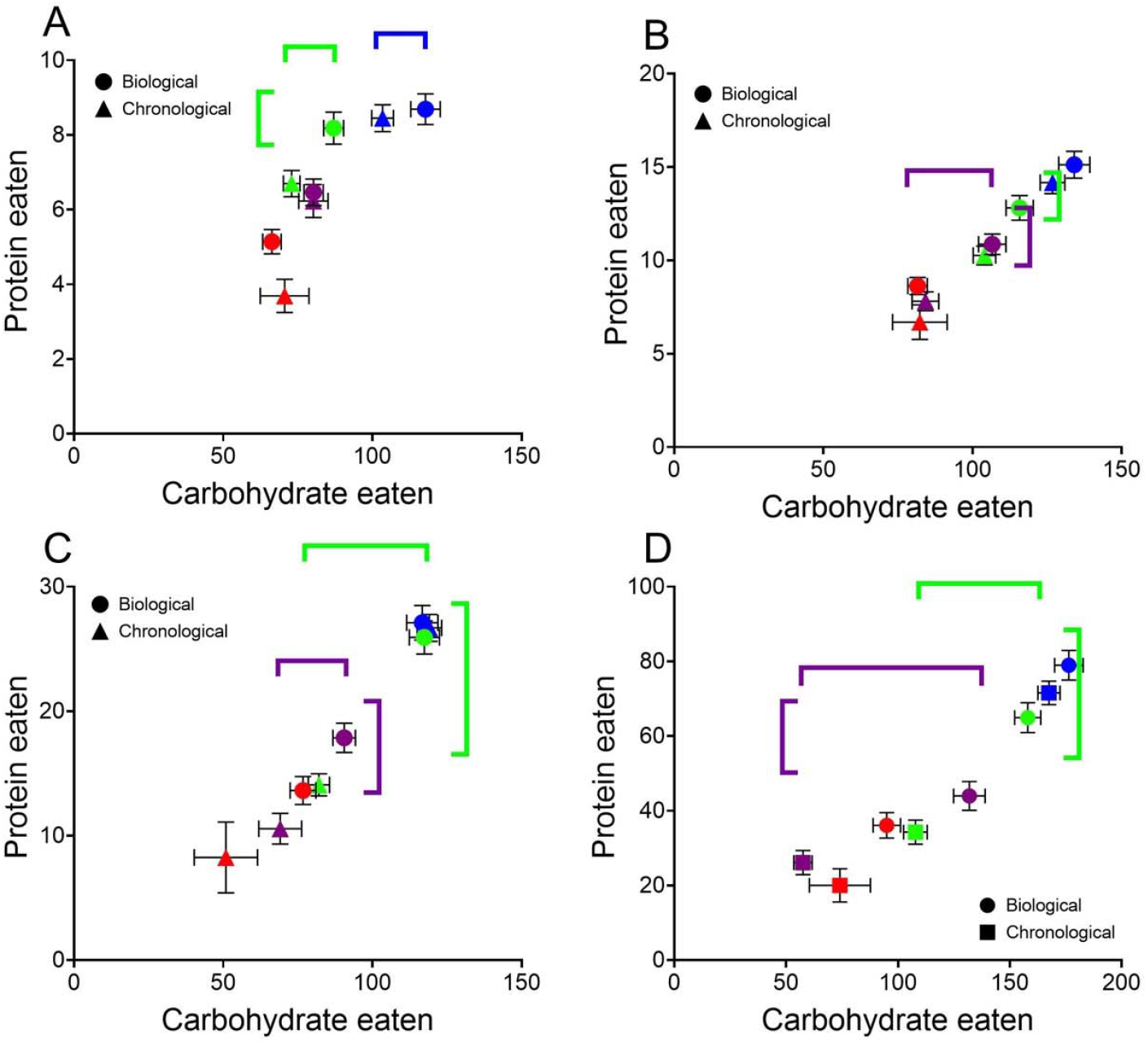
Amounts of protein and carbohydrate consumed by flies at different quartiles of biological and chronological age. Values are represented as mean ± SEM for 10-130 flies fed 15% sucrose and 1.5% (**A**), 2.5% (**B**), 5% (**C**) and 15% (**D**) autolyzed yeast. Horizontal asterisks show significant differences in carbohydrates eaten by flies of the same quartiles and horizontal ones differences in eaten protein. Circles and triangles represent values for quartiles of biological and chronological ages, respectively. Blue, green, violet, and red symbols show values for Q1, Q2, Q3 and Q4, respectively.

Food consumption in female flies is related to nutrient usage in egg production (Mirth et al., 2019). Moreover, egg production induces elevated post-mating feeding in both mated and virgin flies (Barnes et al., 2008). ‘Sex peptide’ (SP) stimulates the female post-copulatory feeding rate (Carvalho et al. 2006). We found the reproductive response to aging in *Drosophila*. Reduced egg laying during aging might predict lifetime reproductive fitness regardless of the diet. Reproductive behavior in *Drosophila* is closely linked to lifespan (Koliada et al., 2020). High reproductive output typically comes at an energetic cost, often resulting in a trade-off between reproduction and longevity (Strilbytska et al., 2024). As *Drosophila* age, reproductive performance typically declines (Churchill et al., 2019; Ruhmann et al., 2018). The rate of this decline can serve as an indicator of biological age, potentially allowing for predictions about lifespan. However, egg-laying activity depends on dietary yeast: tends to increase with yeast concentration, and become stable or decline at extremely high yeast concentrations (McCracken et al., 2020).

Age-induced shifts in physiological parameters observed in our study might be caused by some molecular alterations within the *Drosophila* body during aging. It was previously shown that aging phenotype and longevity may be developmentally programmed (Vaiserman et al., 2018). According to the free radical theory of aging (Harman, 1956), organismal aging is caused by the irreversible accumulation of oxidatively damaged molecules (Lennicke and Cochemé, 2020). Disrupted proteostasis leading to elevated total body protein concentration is considered to be a predictor of aging in *Drosophila* (Yu and Hyun, 2021).

Researchers identified multiple correlations of physiological traits at some time point throughout the chronological age. However, changes in macronutrient intake and reproductive capacity often align more closely with biological age rather than chronological age. For example, the study by Cabrera and colleagues (2020) described the physiological effects of time-restricted feeding on health and aging in *Drosophila*. Lifespan curves demonstrated lifespan extensions at a range between ∼10% and ∼50% in TRF-exposed flies (Cabrera et al., 2020). Moreover, the authors assessed food consumption after 14 or 32 days of TRF and indicated no decrease in food consumption in contrast to dietary restriction. However, it might be assumed that the same amounts of food consumed by TRF-exposed flies and control flies may be caused by the improvement of physiological age caused by the TRF regime. Similarly, the study by Trindade de Paula and colleagues (2016) demonstrated decreased lifespan in flies fed with food supplemented with 10% and 20% of coconut oil. Lifespan shortening in these flies is associated with impaired locomotor ability that was analyzed by negative geotaxis after seven days of exposure to a coconut oil-enriched diet (Trindade de Paula et al., 2016). The reproductive ability of females at different time points throughout chronological age was assessed in the study of Barnes and colleagues (2008). The egg-laying rate against time in days for females exposed to the fertile high-cost differed significantly from the low-cost mating regime (Barnes et al., 2008). Survivorship curves demonstrated the difference in shapes between sterile *ovo*^*D1*^ females exposed to high and low-cost mating regimes (Barnes et al., 2008) suggesting the shift in conformity of biological versus chronological age. Age-dependent motor deficits in *DJ-1* mutant flies may be caused by the higher aging rate in *DJ-1* that was shown in the study of Lavara-Culebras and Paricio (2007). Our current study suggests that the experimental plan for the determination of physiological parameters in *Drosophila* at certain time points should be based on biological age, rather than chronological age. This will allow the researchers to obtain more reliable data, taking into account individual characteristics and the rate of organismal aging.

We found that appetite and reproduction highly correlated with chronological age and survival, as well as with age-related loss of function. These markers decreased with the chronological age of the flies in all dietary regimes and could potentially considered as biomarkers of aging or to predict the aging rate. Flies with preserved reproductive function and balanced nutrient intake tend to exhibit traits associated with delayed biological aging and extended lifespan. Food consumption and fecundity rate are important functional parameters that vary with age and might be used for biological age determination. Understanding the relationship between reproduction, macronutrient consumption, and biological age offers the potential for developing predictive models for lifespan in *Drosophila*.

## 6 Author Contributions

OL: Conceptualization, Data curation, Funding acquisition, Methodology, Writing–original draft, review and editing. OS: Writing–original draft and editing. US, PP and OK: Investigation.

## 8 Acknowledgments

OS acknowledges support from a Virtual Ukraine Institute for Advanced Study (VUIAS) Fellowship program.

